# Gene regulatory network rewiring by an adaptively evolving microRNA cluster in *Drosophila*

**DOI:** 10.1101/227314

**Authors:** Yang Lyu, Zhongqi Liufu, Juan Xiao, Yuxin Chen, Chung-I Wu, Tian Tang

## Abstract

New miRNAs are evolutionarily important but their impact on existing biological networks remains unclear. We report the evolution of a microRNA cluster, *mir-972C*, that arose *de novo* and the subsequently rewired gene regulatory networks in *Drosophila*. Molecular evolution analyses revealed that *mir-972C* originated in the common ancestor of *Drosophila* where it comprises five old miRNAs. It subsequently recruited five new members in the *melanogaster* subgroup after conservative evolution for at least 50 million years. Population genetics analyses reveal that young and old *mir-972C* miRNAs evolved rapidly under positive selection in both seed and non-seed regions. Combining target prediction and cell transfection experiments, we find that sequence changes in individual *mir-972C* members resulted in extensive gene regulatory network divergence among *D. melanogaster, D. simulans*, and *D. virilis*, whereas the target pool of the cluster as a whole remains relatively conserved. Our results suggest that clustering of young and old miRNAs at the same locus broadens target repertoires, resulting in the gain of new targets without losing many old ones. This may facilitate the establishment of new miRNAs within existing regulatory networks.

## 1. Introduction

Newly-evolved genes constitute at least 10–20% of the genome (1, 2) and are among the most conspicuous mechanisms underlying biological innovation (3, 4). Increasing evidence suggests that a large fraction of these new genes is functionally important (1, 4, 5). Using experimental genetic approaches that repress transcripts (6) or disrupt DNA sequences (1, 7) of new genes, previous studies have identified a number of key primarily testes-expressed protein-coding genes that modulate male production (1, 6), sperm competition (8), courtship (9), and pheromone metabolism (10). The functional importance of testes-expressed genes is also supported by the prominent signatures of positive selection at these loci (11). Despite the evolutionary significance of new genes, we know much less about how these novel components integrate into existing regulatory networks. Transcriptomic and protein-protein interaction studies suggest that the targets of new genes change dramatically even among closely related species (7, 12), yet the underlying mechanisms are unclear.

To understand how novel genes mediate among-species divergence of regulatory networks, we need to predict and measure interactions between these new genes and their partners. This is quite challenging for protein-coding genes, as the assays required to establish regulatory or physical interactions are technically demanding. New microRNAs (miRNAs) offer an excellent opportunity to tackle the problem. miRNAs are a class of ubiquitous post-transcriptional regulators that participate in diverse biological processes in eukaryotes (13). In animals, mature miRNAs (~22nt) bind 3’ UTRs of transcripts through perfect base-paring with the seed region (2–8^th^ nucleotides of the mature sequence) (14), inducing translational repression or mRNA degradation (15). It is well-established that the impact of miRNAs on the transcriptome is broad, as each miRNA has hundreds of targets (16). The effects on individual targets, however, are weak. Even the most abundant miRNAs repress single genes by less than 20% (17, 18).

Using next-generation sequencing techniques, previous studies identified a large cohort of new miRNAs from a number of taxa (19). We have previously reported the rapid birth and death of miRNAs in *Drosophila*, a well-studied model system for genetics and molecular evolution. Over 40% of miRNAs are lineage-specific (20). Among these evolutionarily young miRNAs, 95% have appeared *de novo*, and their seed shares no similarities to existing molecules (20). It appears that these newly-evolved miRNAs introduce a wide array of novel miRNA-mRNA interactions. Like new protein-coding genes, new miRNAs are primarily testes-expressed and exhibit strong signatures of positive selection (20). Working out the mechanisms that underlie co-evolution of new miRNAs and testes transcriptomes will provide key insights into new gene evolution processes. For example, we want to know how novel components originate and are assimilated into biological networks. We are also interested in the critical factors that determine whether a new gene will survive and spread into multiple evolutionary taxa.

To understand how newly-evolved miRNAs shape regulatory networks and transcriptomes, and to identify any selective advantage of such systematic change, we focused on a *de novo* originated, fast-evolving and testes-biased miRNA cluster that includes *mir-972* (we refer to the whole cluster as *mir-972C*). It consists of ten miRNA members and is by far the largest new miRNA cluster in *Drosophila melanogaster* (20). We investigate molecular divergence of *mir-972C* among *Drosophila* lineages and study network rewiring in *D. melanogaster, D. simulans*, and *D. virilis*. We discuss the implications of our results for the possible driving forces underlying the evolution of new miRNA clusters and their long-term fate.

## 2. Material and Methods

### (a) Genomic data used

*mir-972C* sequences and coordinates were obtained from miRBase (mirbase.org, Release 19). Genome sequences were retrieved from UCSC (genome.ucsc.edu). The genome versions used here are: *D. melanogaster*, dm3; *D. simulans*, droSim1; *D. sechellia*, droSec1; *D. yakuba*, droYak2; *D. erecta*, droEre2; *D. ananassae*, droAna3; *D. pseudoobscura*, dp4; *D. virilis*, droVir3; *Anopheles gambiae*, MOZ2; *Apis mellifera*, Amel_2.0. GTF annotation files, 3’UTR sequences. 3’UTR locations were downloaded from flyBase (flybase.org, file version: *D. melanogaster*, r6.17; *D. simulans*, r2.02; *D. virilis*, r.1.06). Small RNA and mRNA testes deep-sequence libraries from *D. melanogaster, D. simulans, D. pseudoobscura*, and *D. virilis* (20–24) were retrieved from the GEO database and listed in Table S1.

### (b) miRNA homolog search and phylogenetic inference

We searched for *mir-972C* sequences in downloaded genomes using BLAT with the threshold E < 0.001. miRNA homologs from different species were aligned using MUSCLE with default parameters. To validate miRNAs in *D. melanogaster, D. simulans*, D. *pseudoobscura*, and *D. virilis*, we first predicted secondary structures and thermostability of precursors using RNAfold v2.0 (25) with default parameters. A proper hairpin with free energy less than 15 kcal /mol was considered as a potential miRNA. To validate the expression of miRNAs, small RNA sequence reads (Table S1) were mapped to miRNA precursors using bowtie (parameters: -n 0 -v 0 -f -k 50 -a -p 10) (26). The *mir-972C* homologs covered by at least 10 reads within mature regions were considered as authentic miRNAs. Maximum parsimony analysis was used to infer the origination of the *mir-972C* members by assuming that a miRNA emerged in the most recent common ancestor of all species bearing verified homologs.

### (c) Molecular evolution and population genetics

Molecular divergence, polymorphism, and neutrality tests were computed using in-house perl scripts using the following datasets and methods. Sequence alignments of the *mir-972C* precursors were used to calculate evolutionary divergence (K) between *D. melanogaster* and four other *Drosophila* species: *D. simulans, D. erecta, D. ananassae*, and *D. virilis. D. pseudoobscura* was not included in this analysis because the whole *mir-972C* region was completely lost in this species. Divergence of protein coding genes was calculated as a reference. The protein coding sequence alignment was downloaded from AAA (Assembly/Alignment/Annotation of 12 *Drosophila* Genomes, rana.lbl.gov/drosophila/alignments.html). The Nei-Gojobori model (27) was used to calculate *Ks* (synonymous substitutions per site) and *Ka* (nonsynonymous substitutions per site) of the protein coding genes, and the Kimura’s two-parameter method (28) was used to calculate divergence among miRNA sequences. *D. melanogaster* polymorphic sites were extracted from 50 genomes of the *Drosophila* Population Genomics Project (DPGP; dpgp.org). The *D. simulans* genome (droSim1) was used as an outgroup to infer ancestral states of segregating sites. Three estimates of nucleotide diversity (*θ*), i.e. *θ_π_* (29), *θ_W_* (30), and *θ_H_* (31), were calculated. The DH test was performed as described in (32).

### (d) Target evolution and functional analyses

We predicted target sites based on complementary sequence matches between 3’UTRs and seed regions. To select testes-expressed genes, we used the published testes RNA-seq data (Table S1) and mapped the reads to the genome using STAR (parameters: --runThreadN 4 --runMode genomeGenerate) (33). Read counts at the gene level were calculated by counting all reads that overlapped any exon for each gene using featureCounts (34) and normalizing to TPM (<u>T</u>ranscripts <u>P</u>er Kilobase <u>M</u>illion). We used mean TPM from multiple replicates for our analyses. Genes with TPM ≥ 1 were considered expressed. The number of targets in common among different species was visualized using BioVenn (35). We used DAVID to perform the Gene Ontology (GO) enrichment test for the predicted targets (DAVID v6.7, david.abcc.ncifcrf.gov, EASE < 0.05, adjusted p-value < 0.05) (36).

### (e) *In vitro* validation of miR-975 targets

To construct the *pUAST-mir-975* plasmid from each species, we amplified *mir-975* from the genomic sequences of *D. melanogaster* (ISO-1), *D. simulans* (simNC48S), and *D. virilis* (V46) and cloned the fragments into the *pUAST* vector (see Table S3 for primers and restriction sites). PCR reactions were carried out using the EX-Taq DNA Polymerase (TAKARA). Cells were transfected to 48-well plates with 100 ng of *ub-GAL4* and 200 ng of conspecific *pUAST-mir-975* or the control vector (*pUAST* only) using Lipofectamine 2000 (Thermo Fisher Scientific, catalog no. 12566014), and were harvested 48 hours after transfection.

Total RNA was extracted from cells using TRIzol (Thermo Fisher Scientific, catalog no. 15596026) for qRT-PCR and RNA-seq assays. Total RNAs were reverse-transcribed into cDNAs using stem-loop reverse transcription and analyzed using the TaqMan qRT-PCR method following the miRNA UPL (Roche Diagnostics) probe assay protocol (37). The 2S RNA was used as the endogenous control (see table S4 for the qRT-PCR primers). cDNA libraries for transcriptome assays were generated from each RNA sample and sequenced using Illumina HiSeq 2000 at the Beijing Genomics Institute (Shenzhen). Reads were mapped to genomes using TopHat (v.1.3.1) with the parameter -r 20 (38). Gene expression was estimated use FPKM (<u>F</u>ragments <u>P</u>er <u>K</u>ilobase per <u>M</u>illion) using Cufflinks (v.2.1.1) with default parameters (39). Differentially expressed genes were estimated using Cuffdiff (v.2.1.1) with default parameters (39). Non-expressed genes (FPKM=0) were removed from further analyses. *miR-975* targets were predicted based on seed match using an in-house perl script.

## 3. Results

### (a) Emerging new members in an old miRNA cluster

The *mir-972C* cluster of *D. melanogaster* comprises ten miRNAs spanning a 10.8-kb region located in the 18C-D band of the X chromosome. Based on genomic proximity among members, it was further divided into three sub-clusters (*mir-972/973/974, mir-4966/975/976/977*, *mir-978/979*), spanning less than 1 kb each, and an orphan miRNA *mir-2499* (Figure 1). The *mir-972C* members most likely originated *de novo*, as no sequence similarity was found either between the cluster members, or between them and other miRNAs that have been characterized in *D. melanogaster* (BLAST search, E < 0.001).

**Figure 1.**
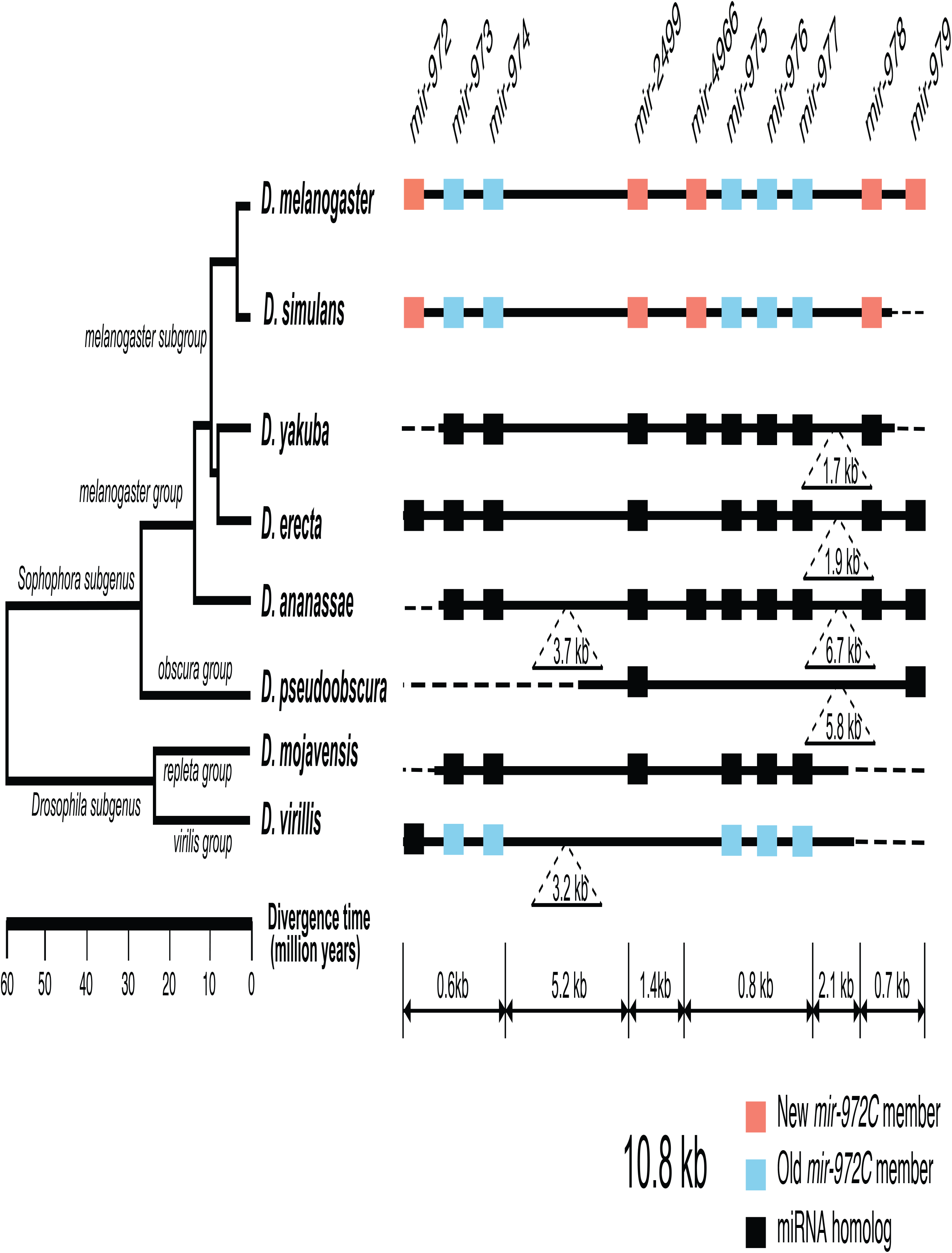
Evolutionary history of the *mir-972* cluster (*mir-972C*) in *Drosophila*. This cluster includes ten miRNAs with distinct seeds. New (red boxes) and old (blue boxes) *mir-972C* members validated through thermostability and deep sequencing are shown. miRNA homologs without expression evidence are colored in black. Deletions and insertions observed in sequence alignments are represented by dashed lines and inverted triangles. The genomic region is not drawn to scale.

To investigate the origin and evolution of *mir-972C*, we searched for orthologs of these miRNA genes in other *Drosophila* species, as well as in *Aedes* (mosquito) and *Apis* (honey bee) which have diverged from *Drosophila* 250 and 300 million years ago, respectively (40, 41). We found homologous sequences in all seven *Drosophila* genomes surveyed (*D. simulans, D. yakuba, D. erecta, D. ananassae, D. pseudoobscura, D. mojavensis*, and *D. virilis*), but failed to detect any homologs in the mosquito or honey bee genomes. This result indicates that *mir-972C* emerged in the common *Drosophila* ancestor between 60 and 250 million years ago. After origination, members of *mir-972C* have undergone rapid birth and death. We only detected *mir-2499* and *mir-979* in the *D. pseudoobscura* genome, suggesting the loss of *mir-972C* members in this lineage. The distribution of individual miRNAs also varies across the remaining species. For example, *mir-973/974/975/976/977* sequences are represented in all seven species, while other miRNAs have been lost in various lineages (Figure 1).

To date the origin time of each miRNA, we first used testes small RNA sequence data to validate that individual *mir-972C* members are expressed in four *Drosophila* species: *D. melanogaster, D. simulans, D. pseudoobscura*, and *D. virilis* (Table S1). A miRNA was considered validated if it (1) showed a predicted hairpin structure with at least moderate thermostability (free energy < -15 kcal/mol) and (2) had more than ten short reads that matched its mature sequence. We found that *mir-973/974/975/976/977* are expressed in *D. melanogaster, D. simulans*, and *D. virilis; mir-972/2499/4966/978* are expressed in *D. melanogaster* and *D. simulans*; and *miR-979* is expressed solely in *D. melanogaster*. No expression of miRNA homologs in *D. pseudoobscura* was detected.

Taken together, these results indicate that *mir-972C* has initially originated in the common ancestor of *Drosophila* and subsequently diverged in different clades. This cluster is largely ancestral in *D. virilis*, lost in *D. pseudoobscura*, and, most interestingly, has been constantly recruiting new members in the *D. melanogaster/D. simulans* branch. Although the cluster originated more than 60 million years ago, the youngest member, *mir-979*, emerged as recently as 4 Myrs ago. Based on the phylogeny, we classified *mir-972C* members into new members that originated after the *Sophophora/Drosophila* split (*mir-972/2499/4966/978/979*) and old members that arose before that event (*mir-973/974/975/976/977*).

### (b) Both new and old *mir-972C* members are evolving rapidly under recent positive selection

We focused on the sequence evolution of *mir-972C* members in *D. melanogaster, D. simulans*, and *D. virilis*, where the mature and miR* sequences were confirmed by deep sequencing. We found the rapid sequence change in both the new and old members (Figure S1). In total, 180 out of 783 sites (23.0%) exhibit substitutions in miRNA precursors. Surprisingly, 28 of the 180 substitutions (15.6%) reside in mature sequences, which are highly conserved in the vast majority of miRNAs. To compare the evolutionary rates between *mir-972C* members and other miRNAs, we calculated the miRNA precursor substitution rates (*K_precursor_*, denoted as *Kp*) between *D. melanogaster* and other *Drosophila* species, normalizing it by genome-wide synonymous site (Ks) substitution rates. Figure 2 shows that the *Kp/Ks* value of *mir-972* members is significantly higher than the entire miRNA population between *D. melanogaster* and closely related species (i.e. *D. simulans* and *D. erecta*, divergence time 4 Myrs and 10 Myrs respectively). However, these differences disappeared once the divergence time was increased to 16 Myrs (i.e. between *D. melanogaster* and *D. ananassae*), or more (i.e. between *D. melanogaster* and *D. virilis*). Consistent with the sequence alignment (Figure S1), both the new and the old members show increased *Kp/Ks* in the last 10 Myrs.

**Figure 2.**
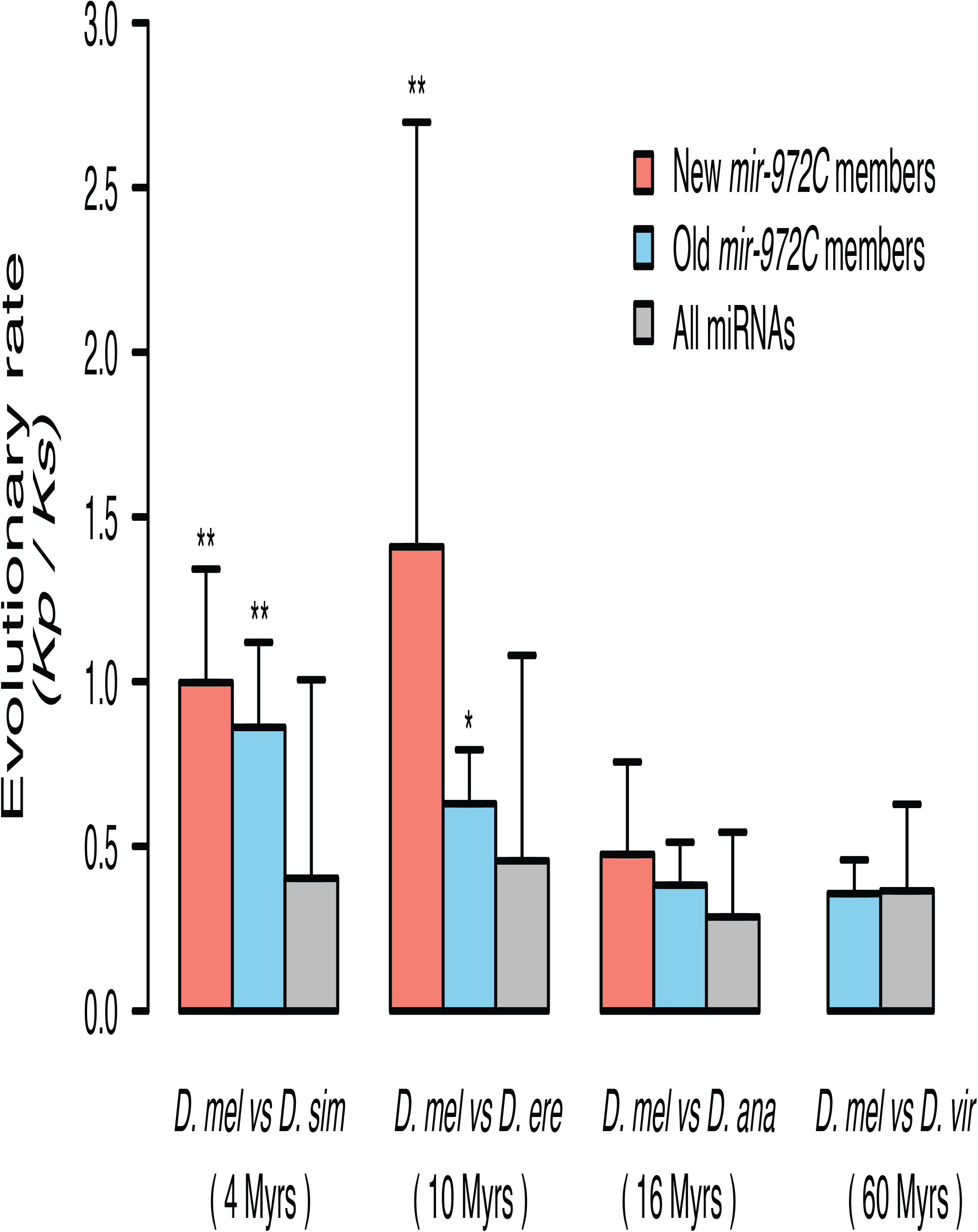
*mir-972C* evolutionary rates compared to all known *Drosophila* miRNAs. Evolutionary rates were measured by *Kp/Ks*. Error bars represent standard deviations (** *p*<0.01, * *p*<0.05, Wilcoxon rank-sum test). Species abbreviations: *D. mel, D. melanogaster; D. sim, D. simulans; D. ere, D. erecta; D. ana, D. ananassae; D. vir, D. virilis*.

Rapid sequence evolution can either be driven by positive selection or relaxation of constraint. In our previous study, the signature of positive selection in *mir-972C* came from an excess of divergence compared to polymorphism of the miRNA precursor sequences in contrast to synonymous sequences genome-wide (20). To further validate positive selection on this cluster we utilized the DH test, a neutrality test based on the allele frequency spectrum in a population (32). The DH test takes advantage of two population allele frequency statistics: Tajima’s D (29) and Fay and Wu’s H (31) to achieve high specificity in detecting positive selection and was shown to be insensitive to demographic changes (32). Table 1 demonstrates that the precursor sequences of five cluster members, *mir-973/974/975/978/979* significantly deviate from the neutral simulations of the DH statistic. *mir-977* also shows a negative D and H, but the p-value is not significant (*p* = 0.06). We were unable to infer the direction of selection operating on the *mir-972/976/2499* cluster due to a lack of polymorphism.

**Table 1.** Polymorphisms and the DH test for individual *mir-972C* members

| locus | $\theta_w (10^{-3})$ | $\theta_\pi (10^{-3})$ | $\theta_H (10^{-3})$ | DH test |
| --- | --- | --- | --- | --- |
| *mir-972* | n/a | n/a | n/a | n/a |
| *mir-973* | 8.0 | 1.6 | 36.9 | **< 1.0E-6 \*\*** |
| *mir-974* | 4.6 | 1.8 | 17.6 | **8.0E-3 \*** |
| *mir-2499* | 14.2 | 11.2 | 3.4 | > 0.5 |
| *mir-4966* | n/a | n/a | n/a | n/a |
| *mir-975* | 3.0 | 0.6 | 0.0 | **2.0E-2 \*** |
| *mir-976* | n/a | n/a | n/a | n/a |
| *mir-977* | 4.7 | 3.4 | 34.5 | 6.0E-2 |
| *mir-978* | 3.8 | 0.8 | 0.0 | **4.0E-3 \*** |
| *mir-979* | 3.1 | 0.6 | 0.0 | **4.0E-3 \*** |

\* <0.05 \*\*<1E-4

Collectively, the evidence supports the inference that evolution of *mir-972C* was driven by positive selection. Members of *mir-972C*, young and old, underwent accelerated evolution only recently. Most importantly, nucleotide changes between species occurred even in the mature sequences which are mostly constrained by purifying selection (42). Such changes in mature miRNAs presumably have profound impact on the downstream regulatory network via shifts in miRNA target repertoires.

### (c) miRNA regulatory network rewiring through seed innovation

To examine the evolution of the regulatory network mediated by miR-972C, we first documented seed changes among *D. melanogaster, D. simulans*, and *D. virilis*. The precursor alignment (Figure S1) reveals two types of seed changes: seed shifting (e.g. *mir-978*) and arm switching (e.g. *mir-975*). We further demonstrate these two events on the phylogenetic tree. As shown in Figure 3a, six of the eight events occurred after the split of *D. melanogaster* and *D. simulans* 4 Myrs ago. Both the new and old *miR-972C* members are involved in this seed innovation. The arm switching of *mir-975* occurred after the split of *D. virilis* and *D. melanogaster/D. simulans* but it is unclear on which branch. *mir-973* is the only member that experienced both seed shifting and arm switching, and its seed is different among all three species (Figure S1).

**Figure 3.**
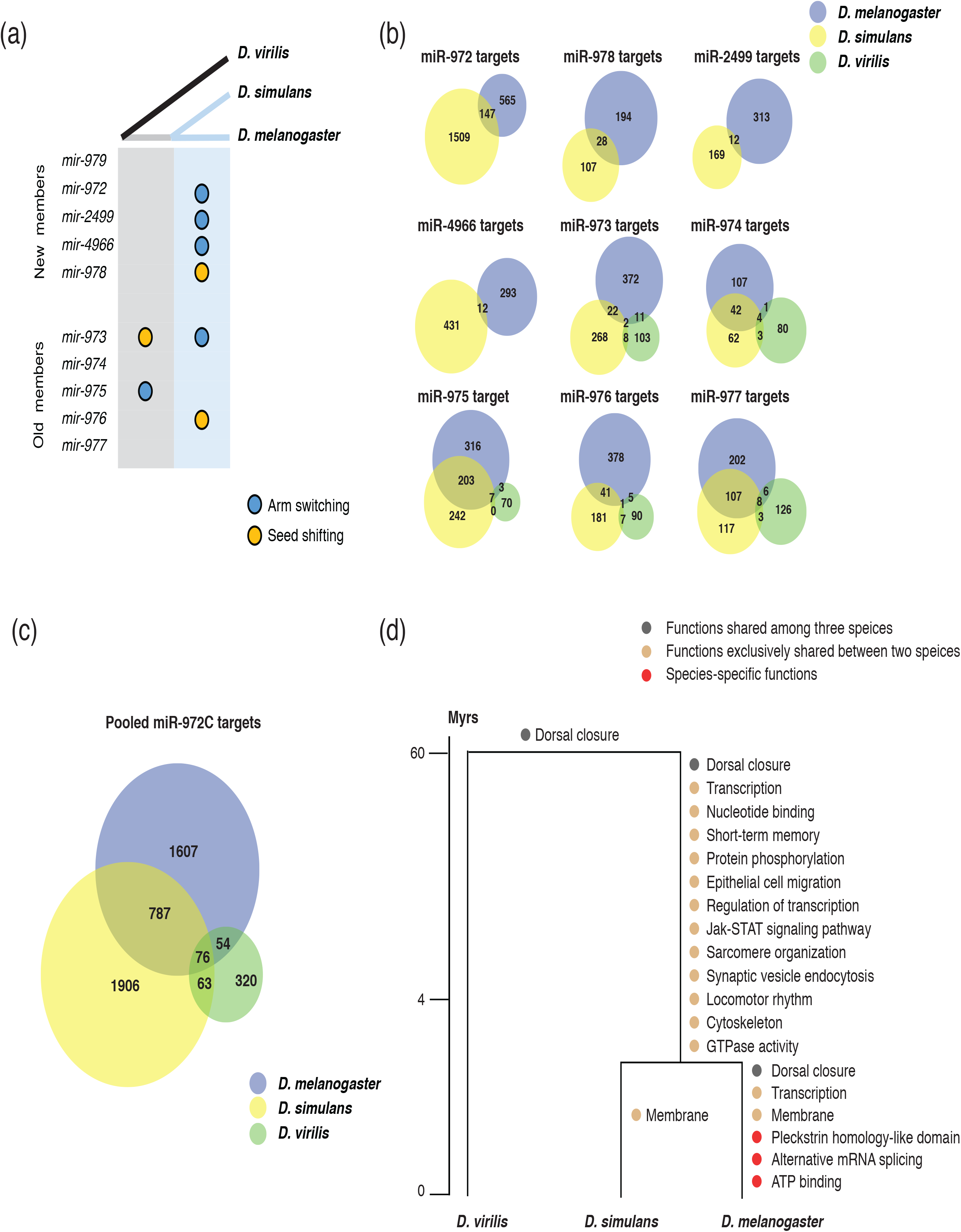
*miR-972C* target and functional evolution. **(a)** *miR-972C* seed innovations. Arm switching (blue circles) and seed shifts (yellow circles) were inferred and denoted along ancestral (grey) and recent (light blue) branches. (**b&c**) Venn diagrams depict the number of shared miRNA targets. **(b)** Individual *miR-972C* member targets. **(c)** Pooled cluster targets. **(d)** Functional evolution of targets. GO categories of shared and lineage-specific targets are indicated on the corresponding evolutionary branches.

Before studying the evolution of *miR-972C* targets, we wanted to make sure that miRNAs and targets are co-expressed in the testes. To this end, RNA-seq reads from the testes of *D. melanogaster, D. simulans*, and *D. virilis* (21, 22) were mapped to their respective genomes and the number of reads within each gene was normalized to TPM (Transcript Per Million). After removing the genes whose expression was not supported by enough reads (TPM < 1), we retained 13,484 genes in *D. melanogaster*, 13,752 in *D. simulans*, and 11,831 in *D. virilis* for further analyses. Using this dataset, we next predicted miRNA targets based on matching of complementary sequences between miRNA seeds and target 3’UTRs for each *miR-972C* member. As shown in Figure 4b, the number of overlapping targets between the *D. melanogaster* / *D. simulans* branch and *D. virilis* is extremely small (<10 for each miRNA), even though the seed sequences remain the same for *mir-974* and *mir-977*. These results suggest that 3’UTR divergence plays a major role in *miR-972C* target evolution.

**Figure 4.**
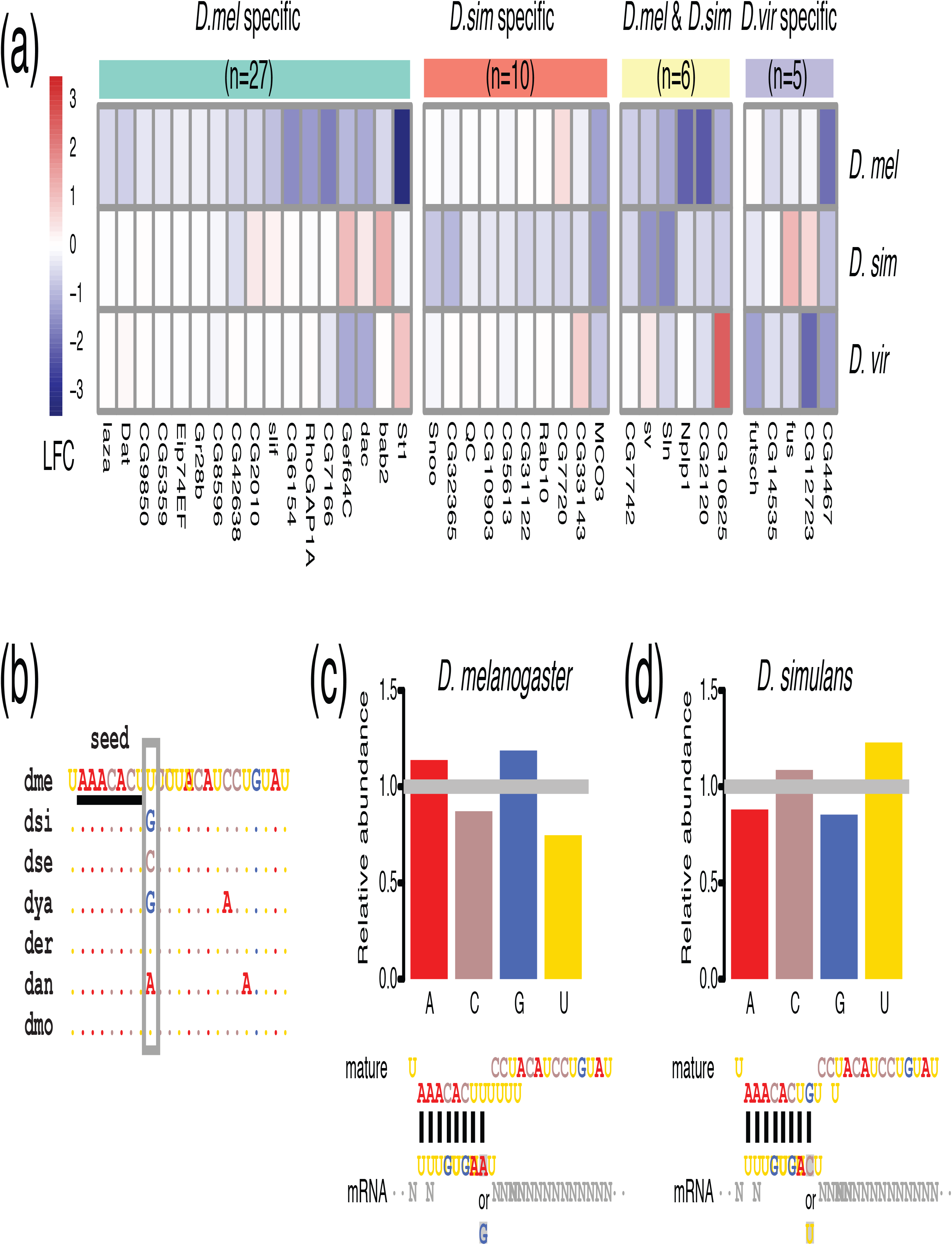
*In vitro* validation of *miR-975* target divergence. **(a)** Heatmap shows the log2(fold change) (LFC) of validated *miR-975* targets in the three species. **(b)** *miR-975* mature sequences from seven *Drosophila* species. They share the same seed (underlined), but their 9^th^ nucleotide (grey box) varies among species. Species abbreviations: dme, *D. melanogaster*; dsi, *D. simulans*; dse, *D. sechellia*; dya, *D. yakuba*; der, *D. erecta*; dan, *D. ananassae*; dmo, *D. mojavensis*. **(c, d)** Bar plot demonstrates that the 3’UTR sites bound to the 9^th^ base of mature sequences are enriched for purines (A or G) in *D. melanogaster* **(c)**, but are enriched for pyrimidines (C or U) in *D. simulans* **(d)**.

We took a closer look at the target divergence between *D. melanogaster* and *D. simulans*. While the *mir-974/975/977* seeds are identical between these species, the proportion of overlapping targets ranges from 21.0% to 27.2% (Figure 3b). Both seed shifting and arm switching significantly reduced the overlap: only 1.6-6.6% of targets are shared between the two species after arm switching (*mir-972/2499/4966/973*), and 6.8-8.5% are shared after seed shifting (*mir-978/976*) (Figure 3b). Although the number of overlapping targets between *D. melanogaster* and *D. simulans* was small for each miRNA after seed changes, the overlap in targets for the entire cluster (19.2%) was largely comparable with that of the miRNAs with identical seeds (Figure 3c). This is likely because a 3’UTR targeted by a *miR-972C* member in one species can be targeted by a different component in another. These observations support the idea that although the targets of each miRNA evolve rapidly, the entire miRNA cluster can keep a relatively conserved target pool.

To understand the functional consequences of *mir-972C* target evolution, we examined the Gene Ontology (GO) enrichment of the predicted targets on each evolutionary branch using DAVID (36) (see Material and Methods and Table S2). Common targets that are shared among three species are enriched in “dorsal closure” (*p* = 7.49E-04), indicating the ancestral function of this cluster (Figure 3d). After the split of *D. virilis* and the *D. melanogaster/D. simulans* branch, the latter two species continued to gain new targets that are involved in “dorsal closure” (*p* = 4.14E-03), suggesting a reinforcement of ancestral functions. We observed a burst of new GO categories on the *D. melanogaster/D. simulans* branch and also in *D. melanogaster*, consistent with the increase in target number in this lineage (Figure 3d). Interestingly, both species acquired targets involved in “membrane” functions independently (*p* = 4.73E-02 in *D. melanogaster* and *p* = 2.02E-02 in *D. simulans*), which may suggest convergent evolution. These results are consistent with the expectation that *miR-972C* target evolution resulted in the acquisition of novel functions without eliminating their ancestral roles.

### (d) Seed and non-seed mutations contribute to regulatory network divergence measured *in vitro*

*mir-975* has undergone substitutions in both seed and non-seed regions and thus offers a great opportunity to compare the effects of the various miRNA functional regions on target expression. The *miR-975* seed in *D. virilis* is completely different from those in *D. melanogaster* and *D. simulans* as a result of arm switching (Figure S1). Furthermore, there is a single nucleotide substitution right after the seed region in the mature miR-975 sequences between *D. melanogaster* and *D. simulans* (Figure S1).

To understand the impact of *mir-975* sequence evolution on target expression, we overexpressed the conspecific *mir-975* sequences in cells derived from *D. melanogaster* (S2), *D. simulans* (ML-82-19a), and *D. virilis* (WR-Dv-1) and monitored the expression changes of both *mir-975* itself and the transcriptome as a whole. Quantitative PCR confirmed that *miR-975* was only expressed in cells transfected with the *pUAST-mir-975* vector and not in cells transfected with the control empty *pUAST* vector (Figure S2). When *mir-975* was overexpressed, predicted targets were significantly downregulated compared to the background transcriptomes of the *D. melanogaster* and *D. simulans* cells (Figure S3a and b, Kolmogorov-Smirnov test, both *p* < 0.05). The repression magnitude was small, consistent with the typically-observed weak miRNA-mediated repression (43). Repression of targets is not significant in *D. virilis* cells (Kolmogorov-Smirnov test, *p* = 0.30, Figure S3c), probably because only a few targets are predicted in this species (n=65).

Using 1.2-fold repression as a cut-off (44), we found 33 targets that were downregulated in *D. melanogaster*, 16 in *D. simulans*, and five in *D. virilis* (Figure 4a). As expected, none of these targets were shared between *D. virilis* and the other two species, indicating that arm switching has changed the target pool completely. Between *D. melanogaster* and *D. simulans*, only 14.0% of the downregulated targets overlapped, consistent with expectation (Figure 3b). Interestingly, target sites complementary to the ninth base of the *miR-975* mature sequence were enriched for purines (A or G) in *D. melanogaster* (*p* < 0.05, Fisher’s exact test, Figure 4c) but enriched for pyrimidines (C or U) in *D. simulans* (*p* < 0.05, Fisher’s exact test, Figure 4d), consistent with the transversion (G->U) change in the mature sequences between these species (Figure 4b). Taken together, our *in vitro* experiments demonstrate that both seed and non-seed changes in *mir-975* contribute to the divergence of miRNA regulatory networks between *Drosophila* species.

## 4. Discussion

New genes continuously contribute to genetic novelty and offer a unique opportunity to understand the evolution of gene regulatory networks (7, 12). As key players in gene regulation, miRNAs regulate their targets weakly and broadly in animals. However, it remains unclear how new miRNAs integrate into the existing regulatory networks (20, 45). Some controversy even surrounds the assertion that new miRNAs have biological functions at all (46). Here we show that the adaptive evolution of the *mir-972C* cluster is accompanied by dramatic evolution of target repertoires between distantly and closely related *Drosophila* species. Changes of both seed and non-seed regions contribute to the target pool evolution. While the rewiring of the *mir-972C* regulatory network has resulted in the acquisition of new targets that represent novel functions in specific lineages, the vast majority of old targets are conserved when we consider the cluster as a whole. These results shed new light on the formation and evolution of new genes in general.

Our results suggest that clustering of new miRNAs may be beneficial to their establishment after emergence (47). Co-expression of miRNAs in a cluster allows them to greatly expand their target pools while functioning coordinately as a unit. Indeed, while only 0.9% to 4.1% of the genome is potentially targeted by individual members of *miR-972C* in *D. melanogaster* (Figure 3b), 14.4% can be influenced by the whole cluster together. Recent studies have shown that large miRNA target pools are evolutionarily beneficial in maintaining stability of gene expression (43). Consistent with this notion, a significant proportion of the *miR-972C* target pool remains unchanged (e.g., 19.2% of the targets are conserved between *D. melanogaster* and *D. simulans*, Figure 3c), even though arm switching and seed shifting occurred frequently between species. The reinforcement of ancestral functional categories of *miR-972C* targets (Figure 3d), on the other hand, also suggests this miRNA cluster continues to recruit additional targets either through the evolution of existing miRNAs or the birth of new hairpins. Such processes also bring novel functions.

The fast-evolving interactions with the transcriptome imply that these miRNAs had never been deeply integrated into the existing gene regulatory networks, calling into question the prospects of long-term survival of these novel miRNAs. A good case in point is the *mir-310/311/312/313* cluster (*mir-310C*) in *Drosophila*, another rejuvenated miRNA cluster of about the same age as *mir-972C* (20). *mir-310C* is known to affect egg morphology, hatchability, and male fertility (48). Redundant and incoherent regulation of multiple phenotypes by *mir-310C* suggest that miRNAs play a role in stability control (48). It is thus not surprising that the miRNA-target interactions could be readily lost. Unlike *mir-310C* that was duplicated from *mir-92a/b, mir-972C* seems to have appeared from non-functional sequence and gained testes expression. Such tissue specificity may make gene loss more likely by limiting pleiotropic effects. Thus, the disappearance of the entire *mir-972* cluster in the *D. pseudoobscura* lineage is not surprising from this perspective.

It should be noted that as a testes-biased miRNA cluster, the fast-evolving *mir-972C* may be associated with rapid turnover of cellular environments in this tissue. It is well established that the testis is the most rapidly evolving tissue due to intense selective pressures associated with sperm competition, reproductive isolation, and sexual conflict (4). Previous investigations in many taxa have demonstrated that male-biased genes evolve relatively quickly at both the sequence and expression level (49, 50). The changes of chromatin states during spermatogenesis allow aberrant transcription which makes testes a hotspot for new gene origination (4). This cellular environment may boost the evolutionary rate of genes with which it has coevolved, including miRNAs (47, 51). Interestingly, *mir-972C* targets did not show GO enrichment in male functions, despite this cluster’s preferential testes expression (Figure 3d). Why would *mir-972C* then be beneficial to the male reproductive system? One plausible explanation is the high complexity of the testes transcriptome (52) that requires substantial regulation to stabilize the system (45). *mir-972C* would be an excellent candidate to do so as it is highly abundant and broadly tied to the testes transcriptome.

## Data accessibility

The raw data are archived in the GEO data repository with accession number GSE107390.

## Author’s contributions

Y.L., C.I.W. and T.T. conceived the study, Y.L., Z.L., J.X. and Y.C. conducted the research, Y.L. and Z.L. analyzed the data, Y.L., Z.L. and T.T. wrote the paper.

## Acknowledgments

This study greatly benefitted from deep sequencing data provided by the generous *Drosophila* community. We appreciate Ming Yang for statistical methods suggestions. We thank Anthony Greenberg and David Paris for valuable language editing.

